# Mapping changes in the spatiotemporal distribution of lumpy skin disease virus

**DOI:** 10.1101/531343

**Authors:** G. Machado, F. Korennoy, J. Alvarez, C. Picasso-Risso, A. Perez, K. VanderWaal

## Abstract

Lumpy skin disease virus (LSDV) is an infectious disease of cattle transmitted by arthropod vectors which results in substantial economic losses due to impact on production efficiency and profitability, and represents an emerging threat to international trade of livestock products and live animals. Since 2015, the disease has spread across many Eastern European countries as well as Russia and Kazakhstan. This rapid expansion highlights the emergent nature of the virus in more temperate regions than those in which LSDV traditionally occurred. The goal of this study was to assess the risk for further LSDV spread in Eurasia through a) analysis of environmental factors conducive for LSDV and b) estimate of the underlying LSDV risk using a fine spatiotemporally explicit Bayesian hierarchical model on LSDV outbreak occurrence information. We used ecological niche modeling to estimate the potential distribution of LSDV outbreaks for 2014-2016. This analysis resulted in a spatial representation of environmental limits where, if introduced, LSDV is expected to efficiently spread. The Bayesian space-time model incorporated both environmental factors and the changing spatiotemporal distribution of the disease to capture the dynamics of disease spread and predict areas in which there is an increased risk of LSDV occurrence. Variables related to the average temperature, precipitation, wind speed, as well as land cover and host densities were found to be important drivers explaining the observed distribution of LSDV in both modeling approaches. Areas of elevated LSDV risks were identified mainly in Russia, Turkey, Serbia, and Bulgaria. Results suggest that prevailing ecological conditions may be compatible with further spread of LSDV in Eurasia, though models should be continually updated to reflect the current epidemiologic conditions. The results presented here advance our understanding of the ecological requirements of LSDV in temperate regions and may help in the design and implementation of prevention and surveillance strategies in the region.

## Introduction

Lumpy Skin Disease is a vector-borne disease of cattle caused by a *Capripoxvirus* (*Poxviridae* family) (Lumpy Skin Disease virus, LSDV), which is clinically characterized by large skin nodules covering the entire body, emaciation, poor milk production, and abortion (Davies, 1982; Woods, 1988). The morbidity rate varies between 10 and 20% while the mortality rate in an infected herd does not usually exceed 5% (OIE, 2018a). The disease results in substantial economic losses due to reductions in productivity and leads to restrictions on international trade of live animals and animal products (Gari et al., 2011; Tuppurainen & Oura, 2012). Since 2012, LSDV has emerged as a major epizootic pathogen given its rapid geographic range expansion out of Africa and into Turkey, Russia, and Eastern Europe, as consequence it was included in the list of notifiable transboundary disease by the World Organization of Animal Health (OIE, 2018b).

In 2006, LSDV was reintroduced into Egypt via imported cattle from East Africa (FAO, 2013) and subsequently emerged throughout the Middle East (FAO, 2013; Tageldin et al., 2014; Al-Salihi & Hassan, 2015). Since 2015 widespread LSDV outbreaks have occurred across several eastern European countries (Russia, Turkey, Greece, Albania, Bulgaria, Montenegro, Serbia, Macedonia) leading to a wave of emergence extending northward and eastward from the Middle East. Before 2013, the northern boundary of the LSDV-affected area was limited to 38 degrees latitude north. In 2014, 2015, and 2016, this boundary shifted from 40 to 42 and 48 degrees north, respectively. Sporadic outbreaks have been detected as far north as 53 degrees north.

The principal route of transmission for LSDV is apparently driven by arthropod vectors, with mosquitoes, ticks and biting flies believed to play a major role (Chihota et al., 2001; Tuppurainen et al., 2011; Tuppurainen & Oura, 2012). Often LSDV transmission is linked to warm and humid weather conditions that are associated with high population densities of biting arthropods (Ochwo et al., 2018). Direct contact between infected and susceptible animals very rarely results in disease transmission (Carn & Kitching, 1995), though it is possible that the infection spreads by contaminated feed and water (Haig, 1957; Al-Salihi & Hassan, 2015). Long-distance disease dispersal can also be facilitated by windborne carriage of vectors and their transportation via vehicles carrying hay and straw (Klausner et al., 2017). The incubation period of LSDV under natural condition was previously reported to range between 1 and 4 weeks (Haig, 1957; Terrestrial Animal Health Code, 2018).

Due to the important role of blood-feeding arthropods in LSDV transmission, its spread and geographical distribution are heavily influenced by the landscape and environmental conditions (Tuppurainen et al., 2013; Abera et al., 2015; Lubinga et al., 2015). Ecological niche models (ENMs) are often used to quantify the potential distribution of species by correlating environmental conditions with known occurrences, providing a tool to assess the influence of the environment and abiotic conditions (e.g., temperature, precipitation and wind speed) on the occurrence and circulation of species and pathogens of interest. ENMs use these associations to characterize the environmental requirements for the disease agent and vector, and subsequently project those relationships into the geographic suitability, (i.e., LSDV) areas that share similar environmental conditions (Elith et al., 2011; Warren & Seifert, 2011; Radosavljevic & Anderson, 2014). ENM has proven to be a useful method to quantify the potential distribution of LSDV in the Middle East (Alkhamis & VanderWaal, 2016). Moreover, there is a need to understand the ecological drivers for LSDV in more temperate regions, such as Russia and Eastern Europe, where LSDV has recently emerged and spread.

ENMs are used to identify potential geographic distributions; however, the presence of a disease or species is not only determined by favorable environmental conditions, but also by the likelihood or risk that the virus could reach a specific location. Bayesian hierarchical statistical approaches allow the inclusion of spatially and temporally explicit features in regression models and capture how disease risk can be related to proximity to areas experiencing outbreaks (Lawson, 2018). Often these approaches have been used to characterize the second order (“local”) underlying variation in risk as either a spatially structured (e.g., spatially autocorrelated) and/or unstructured effect (e.g., random effect). Bayesian hierarchical models provide a framework to link dynamics driving spatial and spatiotemporal distribution against the backdrop of environmental conditions, for example.

Therefore, the purpose of our study was to identify factors associated with LSDV outbreaks from 2014-2016 and explore geographic areas at-risk based on potential ecologically favorable conditions and the spatiotemporal dynamics of disease spread. Our research may shed further insights into the epidemiology of LSDV, and subsequently, these results may contribute to the formulation of surveillance programs that selectively target high-risk cattle areas.

## Material and methods

### Occurrences-data source

We retrieved and curated LSDV outbreak data from Middle Eastern, Central Asian and Eastern European countries in 2014-2016 from the OIE WAHIS interface (OIE WAHID, 2018) (Table 1). Outbreaks were defined as the detection of one or more cases of LSDV per location. Reported outbreaks usually included date of disease detection for groups of cattle herds that were epidemiologically linked and thus could be considered part of the same outbreak event.

**Table 1.** List of OIE reported outbreaks included in the study by region, country and year.

| Country | Region | Number of LSDV outbreaks | Years |
| --- | --- | --- | --- |
| **Iraq** | Middle East | 8 | 2014 |
| **Iran** |  | 6 | 2014 |
| **Turkey** |  | 1294 | 2014 - 2015 |
| **Kazakhstan** | Central Asia | 1 | 2016 |
| **Albania** | Eastern Europe | 218 | 2016 |
| **Armenia** |  | 1 | 2015 |
| **Azerbaijan** |  | 16 | 2014 |
| **Bulgaria** |  | 217 | 2015 - 2016 |
| **Georgia** |  | 4 | 2016 |
| **Greece** |  | 225 | 2015 - 2016 |
| **Macedonia** |  | 356 | 2016 |
| **Montenegro** |  | 63 | 2016 |
| **Russia** |  | 330 | 2015 - 2016 |
| **Serbia** |  | 225 | 2016 |

For each outbreak, the database contained information on the geographical coordinates, date of occurrence, and numbers of susceptible and infected animals in an affected herd. The data were converted into an ESRI shape file for further visualization and analysis.

### Analysis overview

During the first phase of our study, we constructed ENM models in order to identify areas that were potentially suitable for the occurrence of LSDV by assessing associations between outbreak occurrence locations and abiotic (i.e. environmental variables) and interaction with biotic (cattle and small ruminant population densities) variables. The variables that made a contribution of more than 1% via default Maximum Entropy Modeling (MaxEnt) configuration (Phillips et al., 2006), uncorrelated (r<0.6) and known to have biological association to the virus were further considered for ENM calibration. In the second phase of the study, ecological factors selected via the ENM were used in Bayesian spatiotemporal hierarchical models to further elucidate dynamics of spatiotemporal LSDV’s spread.

### Ecological niche model (ENM)

The modern use of ENMs is based on the “BAM” framework (Soberón & Peterson, 2005). A species’ spatial distribution is defined by three components: i) the biotic (B) component is related to availability of hosts, presence and influence of competitors; ii) abiotic (A) environmental conditions, for example temperature and precipitation; and iii) the ability (M) of a given organism to colonize and disperse to biotically and abiotically suitable areas (Romero-Alvarez et al., 2017). The intersection of these three components (i.e. B∩A∩M) determines the area in which a species may occur (Barve et al., 2011). Climatic variables (abiotic conditions) are the main predictors traditionally used in ENM. However, biotic interactions are known to affect pathogen’s spatial distributions; here we incorporated the availability of host species (cattle and small ruminants) alongside abiotic factors in our model of the distribution of LSDV in area M.

### Study area: Model calibration region M

The definition of the study area is an important step for niche modeling given that administrative boundaries are often not epidemiologically meaningful and the choice of M has a strong impact on model predictions (Barve et al., 2011). To define M, we created a regular spatial fishnet grid consisting of square 20 × 20 km cells covering the whole geographic area where LSDV outbreaks occurred (Fig.1). Two different M’s were constructed: a broader M_1_ considered the distance between the centroid of all locations and the farthest outbreak location (238 km), and this distance was used to create a buffer around the outbreaks locations. M_1_ was used as the ENM calibration area assuming that the virus could potentially make long-distance jumps; M_2_ used a 60 km buffer around the outbreak locations, which was based on estimates of spatial spread of LSDV (Mercier et al., 2018). M_2_ was used as a more conservative approach to identify areas with greater risk for LSDV based on factors associated with more localized LSDV spread, as modeled by the Bayesian spatiotemporal models (Fig. 2).

**Figure 1.**
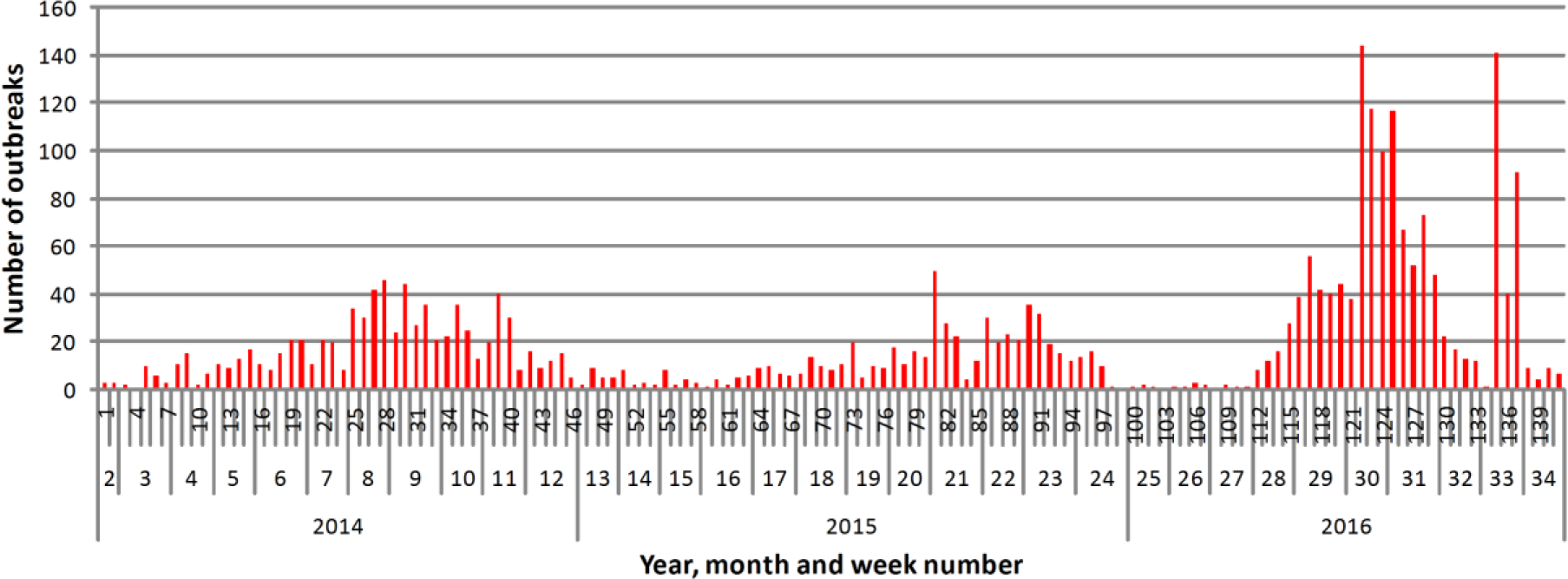
Distribution of the raw number of outbreaks in the study area from February 2014 to October 2016. The x-axis shows the year, month, and week numbered in continuous ascending order.

**Figure 2.**
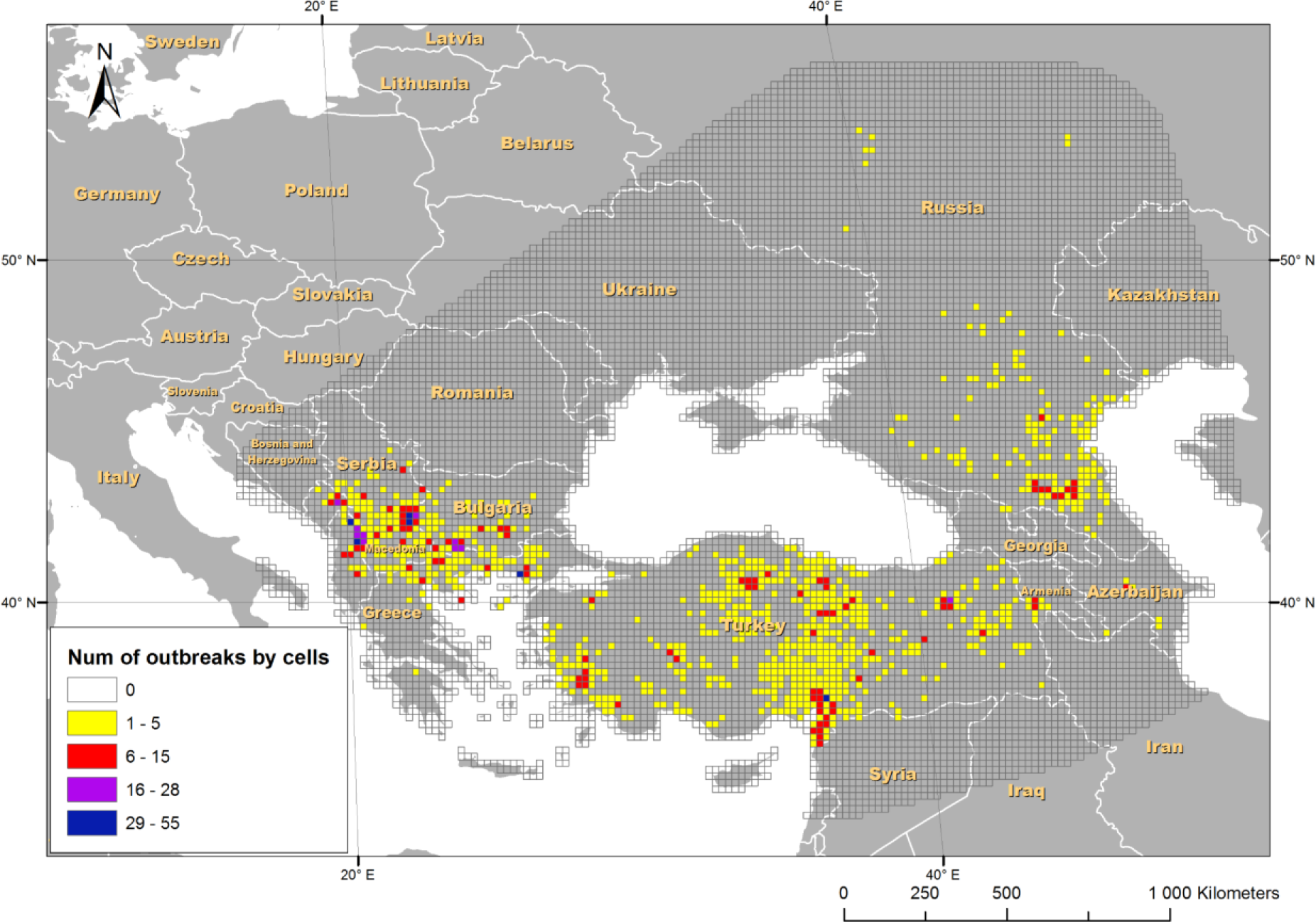
Distribution and frequency of the LSDV outbreaks in the studied area from February 2014 to October 2016. The outbreaks were aggregated into 20×20 km cells-M_1_.

### Independent variables

We included bioclimatic variables (environment), landscape variation (altitude and land cover), and animals density (sheep and cattle) as potential predictors of LSDV suitability. Bioclimatic variables (temperature, precipitation, solar radiation, wind speed, water vapor pressure) were collected from WorldClim2 (Fick &Hijmans, 2017). The original layers consisted of 1×1 km grid cells and allowed for the capture of land surface biophysical patterns and processes over the calendar year (2014 to 2016). WorlClim2 provides access to a monthly average for each variable; we have integrated the average of each variable for each study month in order to provide a better resolution of seasonal effects, which is often ignored by studies of LSDV in which the average values of condition from 1960-1990 are used (Allepuz et al., 2018). For landscape variation, altitude above mean sea level ALT (SRTM based) measured in meters (Jarvis et al., 2008) and land cover types LANDCOVER (MODIS based) (Channan et al., 2014) were used. Cattle and small ruminant population densities (head per square km) were sourced from FAO Gridded Livestock of the World (GLW) (Robinson et al., 2014).

### ENM calibration

Current best practices in ENM include the calibration of several model in which the main goal is to determine the best combination of parameters that represent the phenomenon of interest by finding the best fit to the data (Qiao et al., 2015). We performed ecological niche modeling using Maximum Entropy Modeling (MaxEnt version 3.3.3k) (Phillips et al., 2006) due to its flexibility in fitting a wide range of variable transformations (from linear to hinge), and importantly, its ability to adjust for different parameterizations (regularization values). MaxEnt is an occurrences-background algorithm, which estimates the most uniform probable distribution of the occurrences across a selected calibration region (Phillips et al., 2006) with 10,000 background samples.

Uncorrelated variables which made a contribution of more than 1% in the default MaxEnt configuration were used here to calibrate a wide range of ENMs, including testing 10 regularization coefficients increasing from 0.1 to 5 in 0.5 jumps. The regularization coefficient modulates the fit of the model to the data and model complexity when using several features. If higher values are used, MaxEnt will produce simpler models resulting in broader areas predicted to be suitable for the organism (Merow et al., 2013). Following recommendations (Warren and Seifert, 2011), we have explored a number of feature combinations (L=Linear, Q=Quadratic, P= Product, T=Threshold, and H=Hinge): “L”,“LQ”, “H”, “LQH”, “LQHP”, “LQHPT” “Q”,“LH”,“LT”,“LP”,“QH”,“QT”,“QP”,“HP” (Warren and Seifert, 2011; Merow et al., 2013). These combinations were evaluated using the R package *ENMeval* (Muscarella et al., 2014).

Model fit was assessed initially by Akaike’s Information Criterion (AIC) corrected for small sample size (∆ AICc) (Burnham et al., 2011), so that lower values indicate the least complex model needed to explain the data (Warren & Seifert, 2011). The best regularization coefficient was then used to develop the final model. For the final model, we selected logit values as the output and bootstrap replicates (Elith et al., 2011). Model calibration and evaluation were done via partitioning the occurrences by a k-folds cross-validation approach (Muscarella et al., 2014). Each replicate was based on 80% of occurrences, and 20% for internal evaluation. The threshold to construct the binary suitability map was the minimum logistic value of suitability from 95% of all the occurrences used for model calibration. This threshold takes into consideration an estimate of the likely amount of error among occurrence data and thus removes 5% of occurrences with the lowest suitability values (Peterson et al., 2008).

### Bayesian spatiotemporal hierarchical model

In order to account for the population at-risk, we computed grid cell-specific standardized incidence ratios (SIR), where SIR was defined as 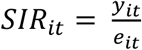 for each spatial cell *i* (*i*=1, …, n=8265) and time period *t* (*t*=0,…, n=33) months. Let *y*_*it*_ be the count of reported outbreaks in cell *i* and month *t*, and *e*_*it*_ the expected number of outbreaks (calculated by multiplying the cattle population in cell *i* for month *t* by the overall number of reported cases of LSDV in M_2_ in month *t*). We used the FAO Gridded Livestock of the World (GLW) (Robinson et al., 2014) cattle population grid to calculate the expected number of cases. Essentially, *SIR*_*it*_ represents the ratio of observed to expected number of outbreaks in a cell (Fig. 2), where the expected count assumes that there is a homogeneous distribution of outbreaks across the study region. A Bayesian hierarchical model was then applied, so that the observed number of outbreaks per cell was assumed to follow a Poisson distribution *y*_*it*_ ~ *Poisson* (*e*_*it*_, *θ*_*it*_) with *e*_*it*_ defined as above and *θ*_*it*_ being the unknown monthly cell-specific relative risk of interest. The model structure allowed for the evaluation of spatially-structured and unstructured variation in LSDV risk, which are known as convolution components (Besag & Mollié, 1991; Lawson, 2013). The structured effect allows for the relative risk within a grid-cell to be dependent on the number of outbreaks occurring within neighboring cells, whereas the unstructured effect captures random variation (Besag & Mollié, 1991; Lawson, 2013).

The first model (LSDV-1) ignored temporal effects and only considered the aggregated number of outbreaks *y*_*i*_, taking into account the spatially structured and unstructured variation in LSDV risk:

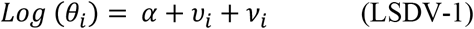

where *α* is the intercept representing the overall level of the relative risk in the study area, and 𝜐_*i*_, 𝜈_*i*_, respectively describe the spatially unstructured and structured variation in LSDV risk across M_2_. A weakly informative Gaussian distribution 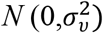 was used as the prior distribution for the spatially unstructured random effect 𝜐_*i*_, while the spatially structured effect 𝜈_*i*_ was modeled with a conditional autoregressive structure as previously described (Knorr-Held & Besag, 1998).

The second model, known as “BYM2”, which included a scaled generalized marginal variance for ν (geometric mean=1), is the Besag-York-Mollié (BYM) model with re-parametrization (Riebler et al., 2016):

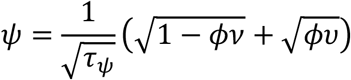

Structured and unstructured temporal effects on the monthly scale *φ*_*t*_ and *γ*_*t*_ were introduced. The structured temporal effect was assigned a random walk type 1 (RW1), where 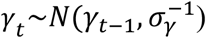, and unstructured temporal effect a 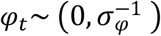 (LSDV-2). Finally, in model (LSDV-3) a space-time interaction term *δ*_*it*_ given by the Kronecker product *γ*_*t* ⊗_ 𝜐_*i*_ was added, and 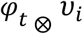 constituted model LSDV-4 such that

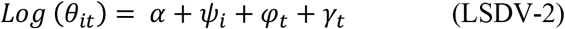

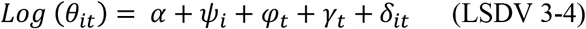

We follow (Simpson et al., 2017; Fuglstad et al., 2017) to construct a join penalizing complexity (PC) prior density, as default we used 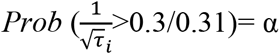 assigned to all the standard deviations of all random effects and for models LSDV 2 to 4 *Prob* (ϕ < 0.5 = 0.7). In addition, we investigated the sensitivity of our results to other less informative priors with larger ranges and strong penalized priors.

We computed the Deviance Information Criterion (DIC) for model selection (Spiegelhalter et al., 2002) and a posterior predictive p-value defined as 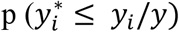, where 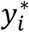 is the predicted value of *y*_*i*_. We considered models with lower values of p-values within the tail deciles as the best fit (Blangiardo & Cameletti, 2015; Baquero & Machado, 2018). Model selection was performed based on the statistically meaningful difference of 3-4 units of DIC (Lesaffre & Lawson, 2012). Finally, the best model was used identity variables associated with LSDV outbreaks. The uncorrelated variables selected through the ENM were considered as potential risk factors candidates. Backwards selection, where only significant covariates were retained, was used to determine the final model. All numeric variables were centered by subtracting their respective mean and scaled by dividing the centered values to the standard deviation prior to any regression analysis.

Finally, we characterized the SIRs predicted by the best model controlling for the spatial risk factors and plotted the posterior mean (which represent the cell-wise RR= SIR). All models were implemented using the Integrated Nested Laplace Approximation (Rue et al., 2009).

### Software

Maxent software package, 3.3.3k version was used for suitability modeling (Phillips et al., 2005). ArcGIS 10.5.1 geographical information system (ESRI, Redlands, CA, USA) with the SDMtoolbox toolkit (Brown, 2014) was used for geographical data processing, geospatial variables preparation and correlation analysis, and for visualization of results. For the descriptive analysis and for data curation, we used R package tidyverse 1.2.1 (Wicham, 2016) and spdep 0.7-4 (Bivand et al., 2015) and INLAOutputs 1.4.11 (Baquero, 2018).

## Results

### Descriptive results

Overall, LSDV outbreaks tended to be seasonal and concentrated in late summer months. However, after the introduction of LSDV to Russia in 2016, there appeared to be an additional pulse of outbreaks in September (Fig. 1). The corresponding geographical distribution of the total number of reported outbreaks for the entire study period is represented in Fig. 2.

### Ecological niche modeling

After the selection of the uncorrelated covariates with contribution above 1%, the following variables were used for ENM model selection: annual mean solar radiation, annual mean wind speed, annual mean water vapor pressure, cattle and sheep population density, altitude, land cover, annual mean maximum temperature and annual mean precipitation. The final MaxEnt ENM was calibrated with a regularization coefficient of 3.1 and LQH features (∆AICc=0). In the final model, variables with the highest percent contribution included annual mean solar radiation (39.1%), annual mean wind speed (33.7%), and annual mean water vapor pressure (10.1%). The results of the final model of the areas potentially suitable for the LSDV outbreaks are shown in Fig 3. Results indicated that Turkey, Greece, Bulgaria, Syria, portions of Iraq and Iran Macedonia, Serbia, Georgia, south of Russia, and Ukraine including still several island and the extreme south of Italy area had consistent suitability for LSDV, while the northern areas including Russia, Kazakhstan and Ukraine were unsuitable (Fig. 3).

**Figure 3.**
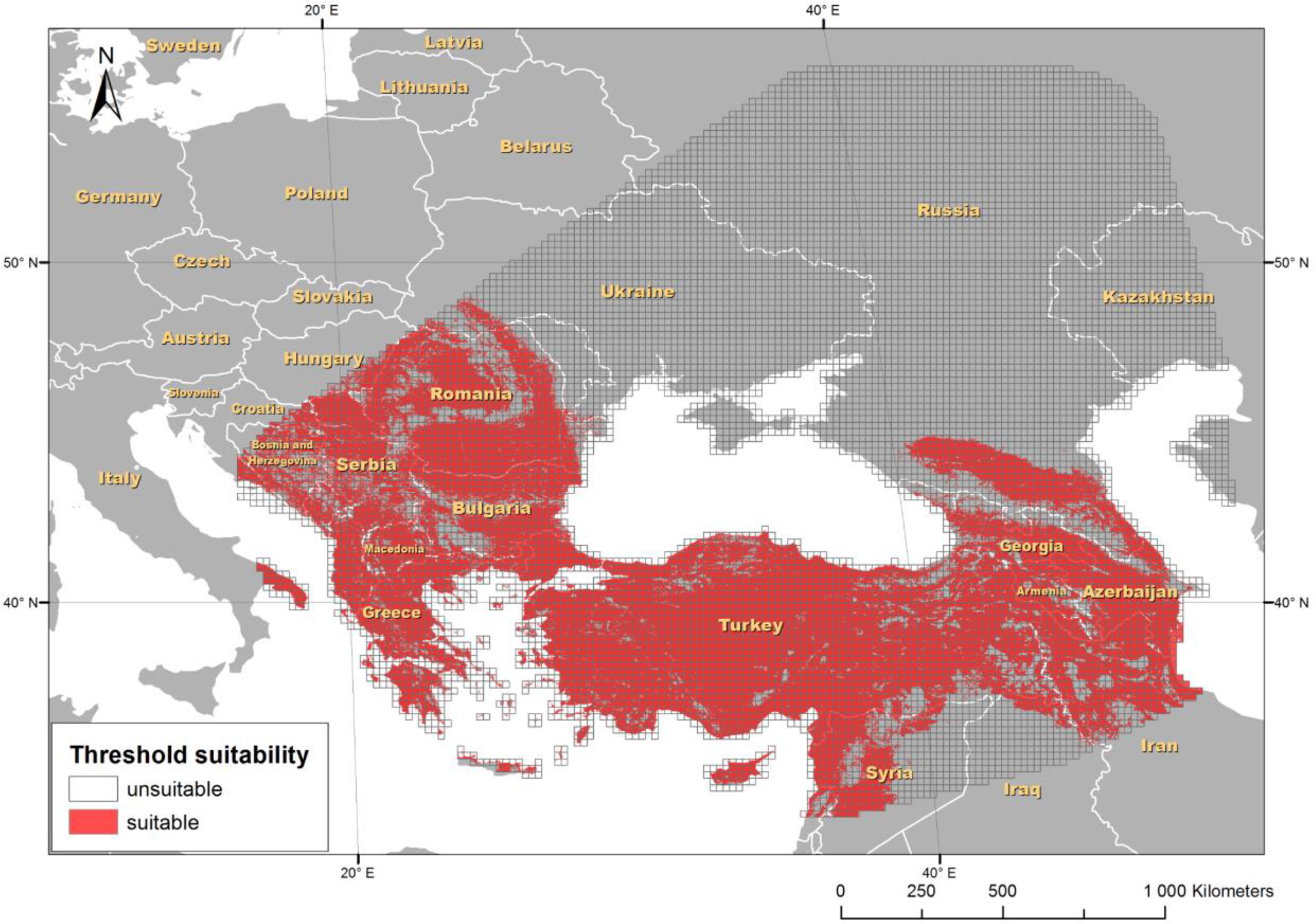
Geographic distribution of LSDV geographic suitability (red) in the study area M_1_.

### Bayesian spatiotemporal hierarchical model

The selected final model included both structured and unstructured spatial effects and a spatiotemporal interaction term among the unstructured effects LSDV-4 (Table 2). Among these effects, the spatiotemporal interaction (**δ)** explained most of the risk variance (56.2%), followed by the spatially structured effect (**ν)** (42.8%), both temporal effects had little contribution (**φ**=0.5% and **γ**=0.4%), and the spatial unstructured effect had a negligible contribution of 0.1% (Table 3).

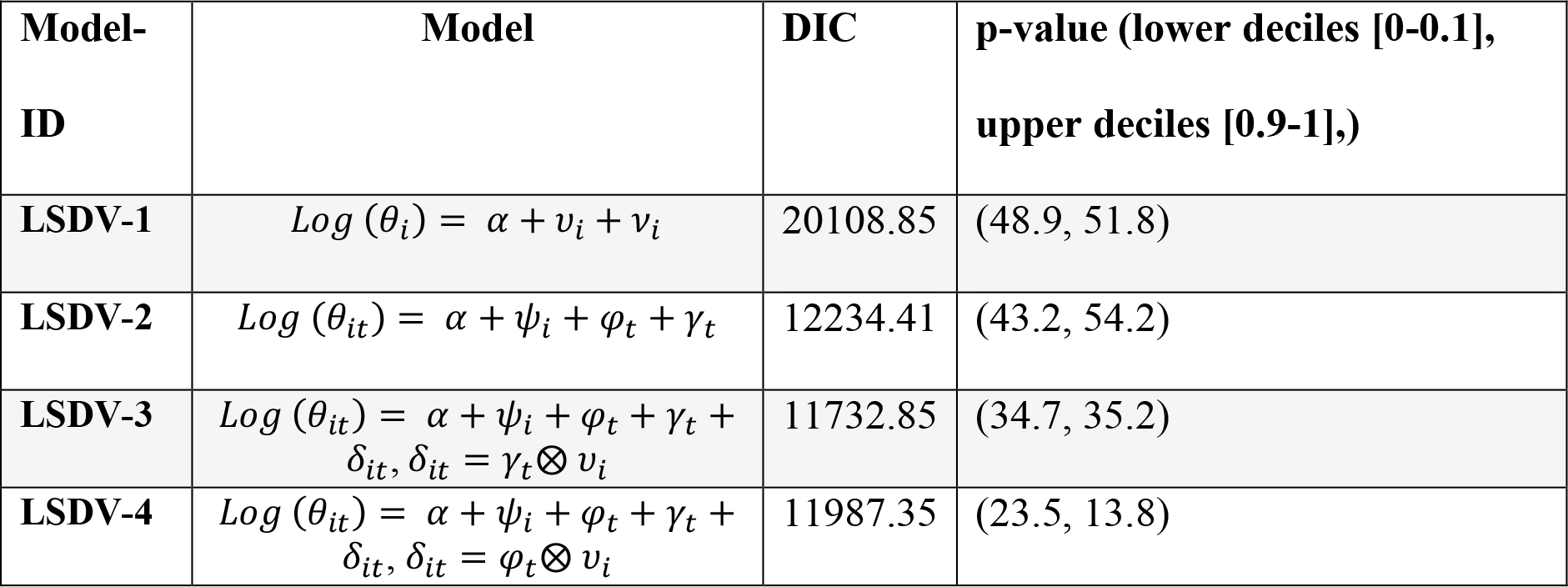

**Table 3.**
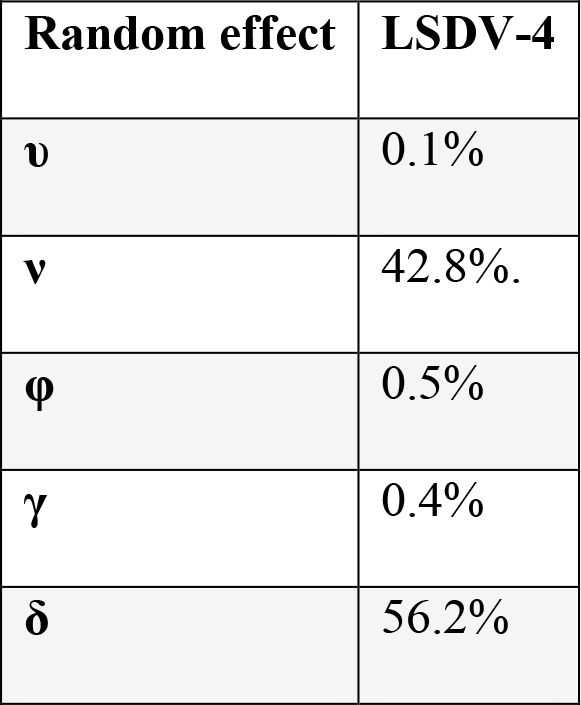
Percent contribution of each spatial and temporal effect to the variance explained by the model: LSDV-4.

The estimated relative risk for LSDV incidence presented a heterogeneous spatial gradient in which values in central Turkey, Bulgaria, Albania and Greece and southeast of Russia were consistently high throughout the study period. Lower risk were found in border areas between southern countries (see Fig.4 for 2016 adjusted RR, see supplement figures 1 and 2 for the monthly estimations of the complete study period).

### Risk factors from the Bayesian spatiotemporal hierarchical model

The final set of variables included in the model Bayesian model selection which met the ENM criteria were average monthly precipitation, average monthly maximum temperature, average monthly solar radiation, average monthly wind speed, average monthly water vapor pressure, land cover and sheep density. In the final LSDV model significant variables were maximum temperature, precipitation and wind. Temperature was the strongest risk factor, with an increase in the incidence of LSDV of 177% for every increase of one standard deviation (Table 4). Likewise, for each standard deviation unit increase in precipitation, number of outbreaks were estimated to increase by 39% (Table 4). Finally, higher wind speeds were negatively associated with LSDV incidence (66% per standard deviation unit increase) (Table 4). While none of the land cover variables were strongly associated with LSDV outbreaks, several land cover classes had posterior distributions that were centered above one (i.e., Arable land, Wooded savannah, and others). Although no single land cover class was strongly associated, the results taken together suggests that land cover may play a role in determining outbreak risk (Table 4).

**Table 4.** Estimated relative risk by model LSDV-4, for LSDV in study area M_2_

| **Model fixed effects** | **RR** | **0.025 quantile** | **0.975 quantile** |
| --- | --- | --- | --- |
| Sheep density | 1.00 | 0.98 | 1.05 |
| Land cover |  |  |  |
| • Coniferous forest | - | - | - |
| • Deciduous broad-leaved forests | 1.33 | 0.52 | 2.78 |
| • Mixed forests | 0.97 | 0.41 | 1.87 |
| • Rare bushes | 0.56 | 0.20 | 1.22 |
| • Wooded savannah | 1.62 | 0.79 | 2.99 |
| • Meadows | 1.33 | 0.73 | 2.34 |
| • Permanent wetlands | 1.04 | 0.23 | 24.25 |
| • Arable land | 1.81 | 0.99 | 3.11 |
| • Urban and built-up areas | 1.48 | 0.69 | 6.50 |
| • Mixed arable land and natural vegetation | 1.32 | 0.53 | 2.70 |
| • Barren lands and wastelands | 84.7 | 0.87 | 23547.1 |
| Vapor pressure | 0.23 | 0.14 | 1.05 |
| Solar | 1.00 | 1.00 | 1.00 |
| **Maximum temperature** | **2.77** | **2.04** | **3.71** |
| **Precipitation** | **1.39** | **1.27** | **1.51** |
| **Wind** | **0.67** | **0.55** | **0.81** |

## Discussion

Our study explored the distribution of LSDV considering two modeling approaches. First, we used ENM to identify additional geographic areas in which, if introduced, may be suitable for LSDV circulation. Second, we used Bayesian hierarchical model to estimate risk by shared information between near neighbors in both space and time to produce new estimates that are more stable and smooth in space and time simultaneously. Finally, we map the combination of environmental suitability and the changing spatiotemporal distribution of the virus. Our integration of these two modeling techniques is novel, and represents an analytical approach to better understand the environmental requirements of a pathogen and identify spatial heterogeneity rates in environmentally heterogeneous regions. In addition, this work provides updated predictions of the environmental suitability and spread of LSDV in more temperate regions.

Our ecological niche model corresponded well to the actual geographic distribution of LSDV, but also highlighted areas where, if introduced, LSDV could efficiently by established and spread. These potentially suitable areas occurred in areas with a monthly median wind speed of 2.4 m s^−1^ (IQR 2.0-2.8), precipitation of ~46.1 mm (IQR 31.5-65.6 mm) with median maximum temperatures of 15.9°C (IQR 8.8-22.8° C), solar radiation of 14633 kJ m^−2^ day^−1^ (IQR 8524-20979 kJ m^−2^ day^−1^) and water vapor pressure of 0.8 kPa (IQR 0.6-1.2 kPa). The suitable areas were between 300 and 1300 meters above sea level (median 782 meters: IQR 299-1306) with sizeable populations of potential hosts such as cattle and sheep (cattle densities median number of 8.3 heads per km^2^ (IQR 4.8-14.78) and sheep 17.1 heads per km^2^ (IQR 6.2-38.4). The model also exclude some areas that were likely unsuitable for LSDV. Because the data used to calibrate our models included outbreaks until October 2016, it was unable to forecast the new cases recently reported in the border with Kazakhstan (Allepuz et al., 2018). Several hypothesis may explain the failure of this model to identify areas where new cases occurred post October 2016. First, we may not be able adequately consider long-distance movement of people and animals (Mercier et al., 2018), nor did we include the distribution of the possible wildlife hosts or vectors richness (Tuppurainen et al., 2018). Consequently, the models generated mainly reflect the areas which had reported cases, an indication of model overfit. This highlights the importance to continually update ENMs for highly dynamic disease situations, particularly for pathogens that are undergoing geographic invasion range expansions which can be compared with the behavior of an invasive species. Further studies should be performed to confirm the capacities of ENM to forecast the potential distribution of LSDV, moreover, models should also include future climate scenarios in order to identify potential shifts in the disease distribution special in areas of translational elevation which may become suitable for the occurrence of potential vectors (Hopp & Foley, 2001; Patz et al., 2005; Gubbins et al., 2018; Tuppurainen et al., 2018b).

The Bayesian spatiotemporal model identified potential patterns of LSDV during the study period and identified potential risk factors. This study is the first attempt to explore nonlinear trends in LSDV epidemic areas. We identify significant spatiotemporal variation in disease risk, which was expected since most infectious diseases have multiple causes that often vary in the space and in time. We captured this variation by ignoring administrative boundaries and instead used a grid cell constructed based in previously studies that estimated the potential distance that LSDV could potential travel (Mercier et al., 2018). The model identified spatiotemporal variation and highlighted the importance of the interaction term between the space and time effects, which was more relevant than the spatial structured risk itself in explaining LSDV outbreaks (Table 3), previous studies have neglected the inclusion of space-time (Allepuz et al., 2018), which is known to produce more accurate models and reveal more spatial and temporal structure (Zehang et al., 2019). The identified hot areas with increased infection risk after controlling for the contribution of maximum average temperature, monthly precipitation and average wind speed were mapped (Fig.4.). Analyzing the spatial risk over several months (i.e., months 6, 7, 8, 18 and 29, see Fig. 4 and supplementary information 1 for RR for all months) showed consistent risk activity at the border of Russia and Kazakhstan in which new cases were recorded in 2017 and 2018 (Allepuz et al., 2018).

**Figure 4.**
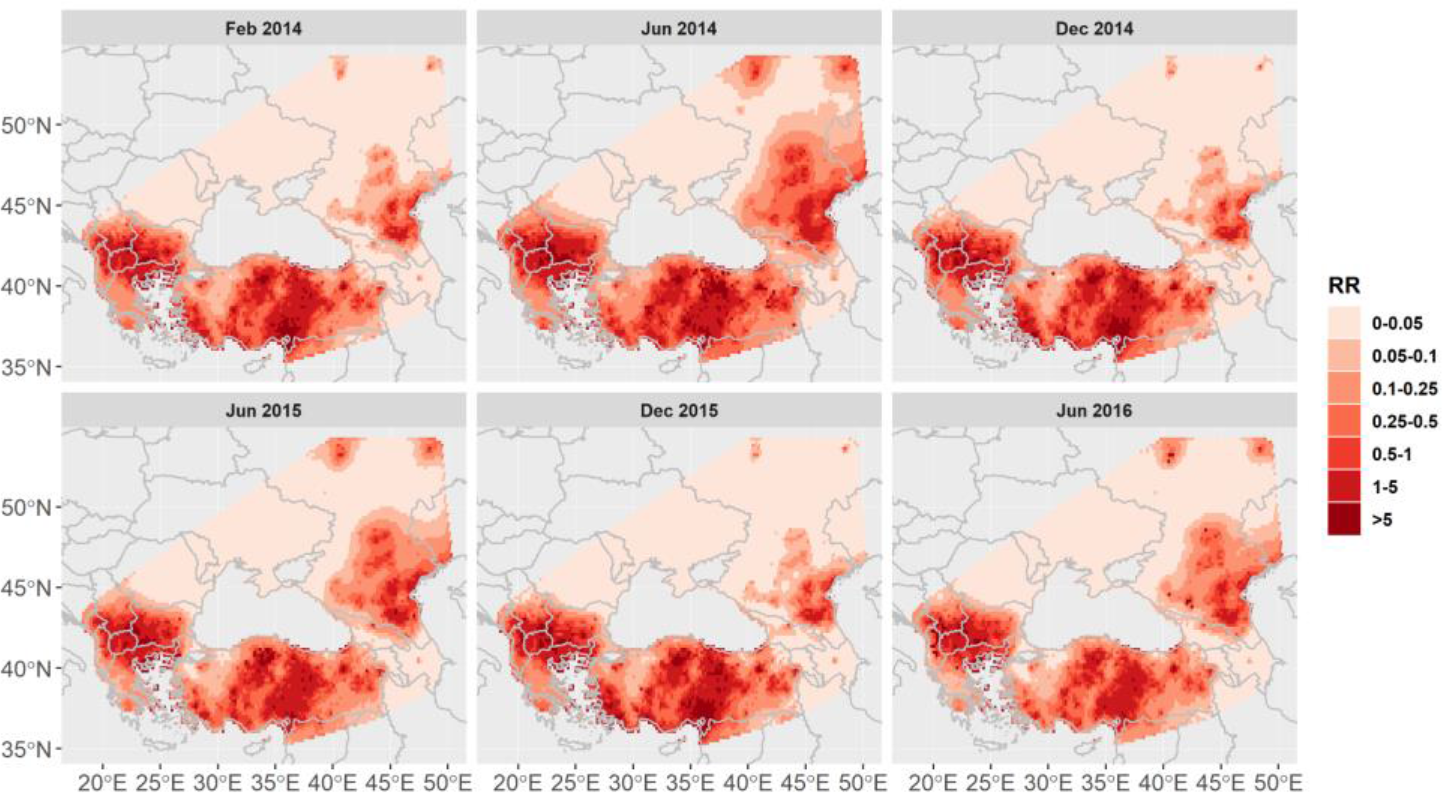
Monthly distribution of the adjusted relative risk (RR) for LSDV in the study area M_2_ according to model LSDV-4 (see supplementary information for RR for all months). The selected months capture the temporal dynamics of the estimated risk. Cell-specific RR values are relative to the mean RR.

The spatiotemporal variability of LSDV was influenced by several independent variables. Risk was positively associated with precipitation and temperature and negatively affected by wind. These findings are in agreement with the majority of the previous research (Alkhamis & VanderWaal, 2016; Tasioudi, et al., 2016; OIE, 2016; Hunter & Wallace, 2001; Eeva, et al., 2018). A contradiction and unresolved topic is the role of wind in the spread of virus or potential vectors, such as *Stomoxys calcitrans*. Correlations have been shown between vector abundance and LSDV circulation (Kahana-Sutin et al., 2017). It is hypothesized that wind could potently transport live virus or infected vectors long-distances, which would explain the large spatial jumps LSDV has been recently shown to make (Klausner et al., 2017). Wind has been reported to be positively associated with disease transmission (EFSA AHAW Panel, 2015; Saegerman et al., 2018), and negatively associated with disease occurrences (Saegerman et al., 2018). We found a negative effect of wind speed, where risk of LSDV is predicted to be reduced when winds are stronger. This is in agreement with a study calculating the risk of LSDV introduction to France, which found that the chances of passive transportation by wind to France would be negligible (Saegerman et al., 2018). It is also possible that, in our models, wind was correlated with some other spatially variable factor that was not captured by any other variables included in our model.

We also identified temperature as increasing the relative risk for LSDV, which has also been described in several other publications (OIE, 2016; Hunter & Wallace, 2001; Allepuz et al., 2018). In a recent report from EFSA that used a very similar approach where the region where LSDV occurred was divided into a 25×25 km grid, it was found that the temperature was a good proxy for disease occurrence in Albania and also in line with the relative abundance of *Stomoxys calcitrans*, a putative vector of LSDV (EFSA AHAW Panel, 2015). Another study also identified higher temperatures associated with increased risk of LSDV outbreaks. This study used a smaller 10×10 km cells, though no biological or epidemiological explanation was provided for the chosen grid size (Allepuz et al., 2018). Precipitation was also associated with increased risk of LSDV in the Middle East (Alkhamis & VanderWaal, 2016).

Our model also suggests that land cover may play a role in determining risk, given that there was a trend for several land cover classes (e.g., Arable land, Wooded Savannahs, and Meadows) to be associated with LSDV outbreaks. For example, arable land had a relative risk of 1.8, but the 95% credibility interval (based on the posterior distribution) suggested that this value could be as low as 0.99 or as high as 3.1. It is possible that the importance of land cover could be better assessed if the 11 classes were condensed to fewer categories. Although no single land cover class was strongly associated with risk in our model, the combined influence of several land cover classes, taken together, suggests that land cover may play a role in determining outbreak risk.

Warm temperatures, wetter conditions, and land cover have been associated with higher activity of insects (Ali et al., 2012). Evidence strongly suggest the involvement of haematophagous arthropod vectors in LSDV transmission among cattle by the mechanical route, but the full vector range of LSDV has not been established (EFSA AHAW Panel, 2015). The importance of different mechanical vectors in the transmission of LSDV is likely to vary in different geographical regions, depending on the environment, temperature, humidity and abundance of the vectors. Further studies are needed to explicitly model the abundance of all possible vectors (EFSA AHAW Panel, 2015, Gubbins et al., 2018). In addition, a limitation of the study was the exclusion of reported cases after October 2016, one reason we decided not to use data from 2017 was the significant reduction in the number of reported cases (Tuppurainen & Oura, 2012; Casal et al., 2018) and we were interested in modeling the disease in its epidemic mode. In addition, we wanted to avoid the influence of the vaccination campaign that were implemented in later 2016 early 2017 in epidemic countries (EFSA, 2018). However, if data were available on the spatiotemporal extent of vaccination, it would be valuable to include in future modeling studies. Finally, although our approach was effective in capturing the spatiotemporal dynamics of LSDV from 2014 through 2016, the model has limited ability to forecast future risk, given that risk is highly linked to the contemporary spatial distribution of cases. This further highlights the need to continually update risk models to reflect dynamic epidemic situations. Our current efforts are on building a pipeline with the capacities for a near-real time risk forecast in the region.

The ENM developed in this study demonstrated the ability to estimate the past distribution of LSDV, which highlighted a positive relationship between risk and water vapor and negative effect of wind and solar radiation. Studies including future climate scenarios are needed and could help to validate the current models for precise estimation of future disease circulation. The spatiotemporal patterns of LSDV occurrences modeled by the Bayesian hierarchical approach were heterogeneous in the region, with temperature and precipitation increasing an area’s relative risk and stronger winds reducing risk. With this study, we identified hotspot areas by using two modeling approaches, demonstrating how these approaches can be integrated to guide disease control and active surveillance efforts.

## Supporting information

Fitted RR spacextime

## Acknowledgements

This project was funded by the Department of Population Health and Pathobiology, College of Veterinary Medicine, North Carolina State University, which provided startup funds for GM; University of Minnesota Academic Health Center Grant-in-Aid, and JA is the recipient of a Ramón y Cajal postdoctoral contract from the Spanish Ministry of Economy, Industry and Competitiveness (MINECO) (RYC-2016-20422).

## Conflict of interest

The authors declare that there are no conflict of interests

## Supplementary information

### Supplementary information 1

Monthly distribution of the adjusted relative risk (RR) for LSDV in the study area M_2_ according to model LSDV-4 (see supplementary information for RR for all months). Temporal dynamics of the estimated risk from February 2014 until October 2016. Cell-specific RR values are relative to the mean RR.

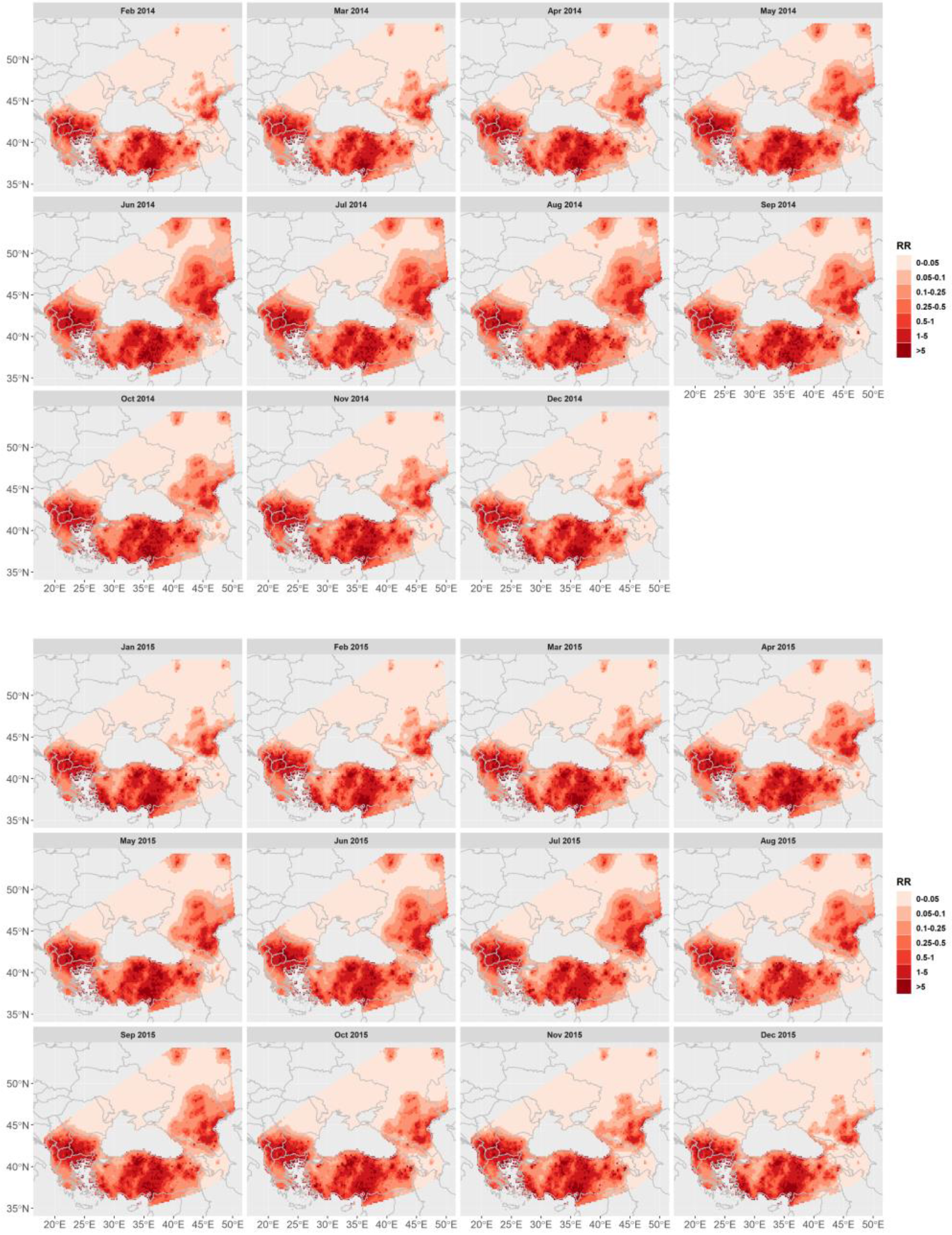

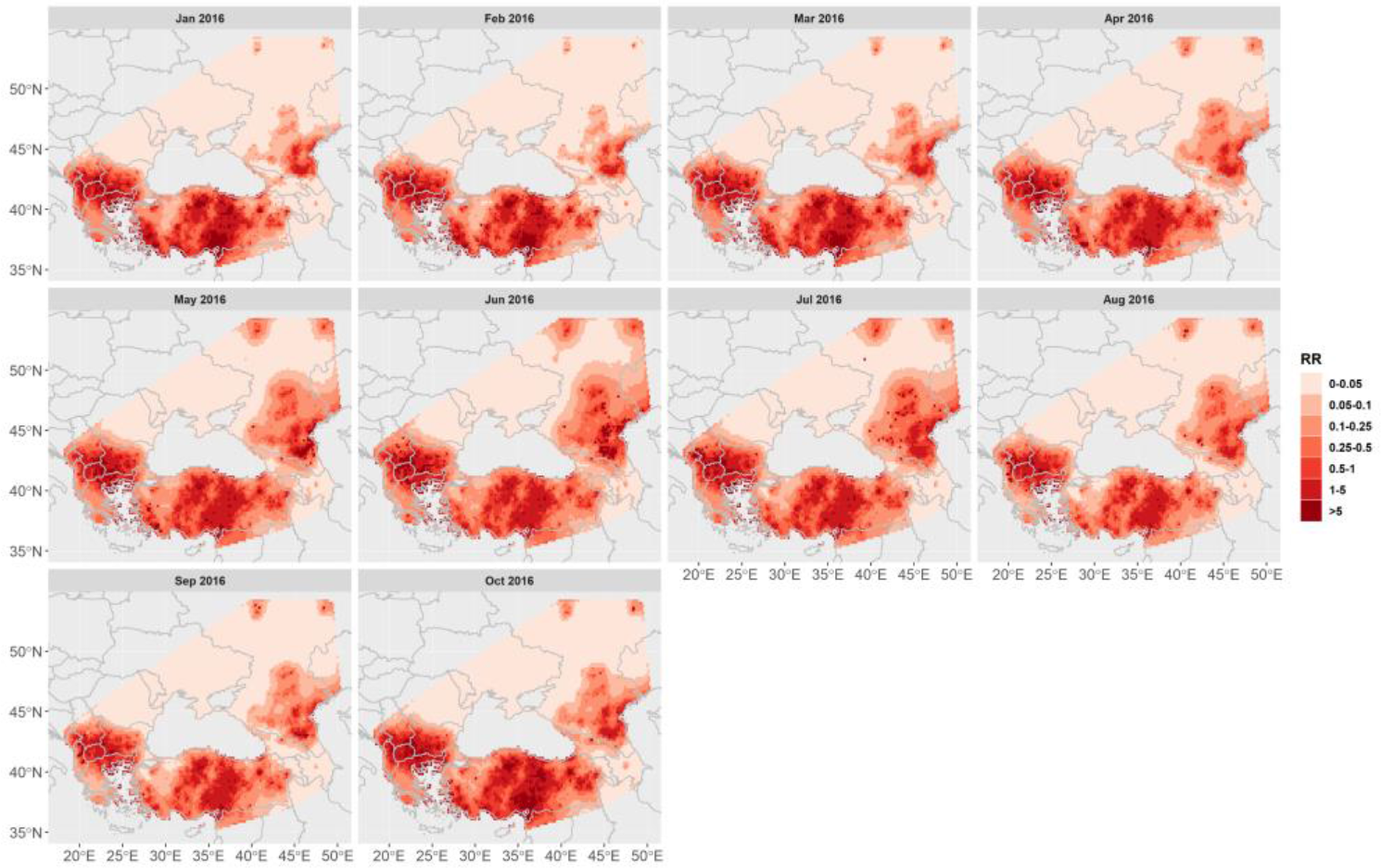

## References

Ali, H., Ali, A.A., Atta, M.S., Cepica, A., 2012. Common, Emerging, Vector-Borne and Infrequent Abortogenic Virus Infections of Cattle. Transbound. Emerg. Dis. https://doi.org/10.1111/j.1865-1682.2011.01240.x

Alkhamis, M.A., VanderWaal, K., 2016. Spatial and Temporal Epidemiology of Lumpy Skin Disease in the Middle East, 2012-2015. Front. Vet. Sci. 3. https://doi.org/10.3389/fvets.2016.00019

Allepuz, A., Casal, J., Beltran-Alcrudo, D. 2018. Spatial analysis of lumpy skin disease (LSD) in Eurasia -Predicting areas at risk for further spread within the region. Transboundary and Emerging Diseases. Dec. 6. https://doi.org/10.1111/tbed.13090. [Epub ahead of print]

Abera, Z., Degefu, H., Gari, G., Kidane, M., 2015. Sero-prevalence of lumpy skin disease in selected districts of West Wollega zone, Ethiopia. BMC Vet. Res. 11. https://doi.org/10.1186/s12917-015-0432-7

Al-Salihi, K.A., Hassan, I.Q., 2015. Lumpy Skin Disease in Iraq: Study of the Disease Emergence. Transbound. Emerg. Dis. 62, 457–462. https://doi.org/10.1111/tbed.12386

Baquero, O.S., Machado, G., 2018. Spatiotemporal dynamics and risk factors for human Leptospirosis in Brazil. Sci. Rep. 8. https://doi.org/10.1038/s41598-018-33381-3

Baquero, O. S. INLAOutputs: process selected outputs from the ‘INLA’ Package. at http://oswaldosantos.github.io/INLAOutputs (2018).

Barve, N., Barve, V., Jiménez-Valverde, A., Lira-Noriega, A., Maher, S.P., Peterson, A.T., Soberón, J., Villalobos, F., 2011. The crucial role of the accessible area in ecological niche modeling and species distribution modeling. Ecol. Modell. 222, 1810–1819. https://doi.org/10.1016/j.ecolmodel.2011.02.011

Besag, J., York, J., Mollié, A., 1991. Bayesian image restoration, with two applications in spatial statistics. Ann. Inst. Stat. Math. 43, 1–20. https://doi.org/10.1007/BF00116466

Bivand, R., Altman, M., Anselin, L., Assunção, R., Berke, O., Bernat, A., Blanchet, G. 2015. Package ‘spdep’. Available at ftp://garr.tucows.com/mirrors/CRAN/web/packages/spdep/spdep.pdf (accessed 17 December 2018)

Blangiardo, M., Cameletti, M. 2015. Spatial and Spatio-temporal Bayesian Models with R - INLA. (Wiley - Jhon Wiley and Sons, Ltd.

Brown, L. J. 2014. SDMtoolbox: a python‐based GIS toolkit for landscape genetic, biogeographic and species distribution model analyses. Methods Ecology Evo. 5 (7), 694–700. https://doi.org/10.1111/2041-210X.12200

Burnham, K.P., Anderson, D.R., Huyvaert, K.P., 2011. AIC model selection and multimodel inference in behavioral ecology: Some background, observations, and comparisons. Behav. Ecol. Sociobiol. https://doi.org/10.1007/s00265-010-1029-6

Carn, V.M., Kitching, R.P., 1995. An investigation of possible routes of transmission of lumpy skin disease virus (Neethling). Epidemiol. Infect. 114, 219–226. https://doi.org/10.1017/S0950268800052067

Casal, J., Allepuz, A., Miteva, A., Pite, L., Tabakovsky, B., Terzievski, D., Alexandrov, T., Beltrán-Alcrudo, D. 2018. Economic cost of lumpy skin disease outbreaks in three Balkan countries: Albania, Bulgaria and the Former Yugoslav Republic of Macedonia (2016-2017). Transboundary and Emerging Diseases. 10 July 2018. https://doi.org/:10.1111/tbed.12926

Channan, S., K. Collins, and W. R. Emanuel. 2014. Global mosaics of the standard MODIS land cover type data. University of Maryland and the Pacific Northwest National Laboratory, College Park, Maryland, USA.

Chihota, C.M., Rennie, L.F., Kitching, R.P., Mellor, P.S., 2001. Mechanical transmission of lumpy skin disease virus by Aedes aegypti (Diptera: Culicidae). Epidemiol. Infect. https://doi.org/10.1017/S0950268801005179

Davies, F.G., 1982. Observations on the epidemiology of lumpy skin disease in Kenya. J. Hyg. (Lond). https://doi.org/10.1017/S002217240006993X

Eeva, S. M.T., Shawn, B., Eyal, K., 2018. Lumpy Skin Disease, Springer, Gewerbestrasse, Switzerland, ISBN 978-3-319-92411-3

EFSA AHAW Panel (EFSA Panel on Animal Health and Welfare), 2015. Scientific Opinion on lumpy skin disease. EFSA Journal. 2015; 13(1):3986, 73 pp. https://doi.org/10.2903/j.efsa.2015.3986

EFSA (European Food Safety Authority), 2018. Scientific report on lumpy skin disease II. Data collection and analysis. EFSA Journal. 2018. 16(2):5176, 33 pp. https://doi.org/10.2903/j.efsa.2018.5176

Elith, J., Phillips, S.J., Hastie, T., Dudík, M., Chee, Y.E., Yates, C.J., 2011. A statistical explanation of MaxEnt for ecologists. Divers. Distrib. https://doi.org/10.1111/j.1472-4642.2010.00725.x

FAO. 2013. Emergence of lumpy skin disease in the Eastern Mediterranean Basin countries. EMPRES WATCH, Vol. 29, November 2013. Rome.

Fuglstad G.-A., Simpson D., Lindgren F., Rue H. 2017. Constructing priors that penalize the complexity of Gaussian random Fields, https://arxiv.org/pdf/1503.00256.pdf

Gari, G., Bonnet, P., Roger, F., Waret-Szkuta, A., 2011. Epidemiological aspects and financial impact of lumpy skin disease in Ethiopia. Prev. Vet. Med. https://doi.org/10.1016/j.prevetmed.2011.07.003

Gubbins, S., Stegeman, A., Klement, E., Pite, L., Broglia, E., Abrahantes, J. C. 2018. Inferences about the transmission of lumpy skin disease virus between herds from outbreaks in Albania in 2016. In press, https://doi.org/10.1016/j.prevetmed.2018.12.008.

Haig, D. A., 1957. Lumpy skin disease. Bull. Epizoot. Dis. Afr. 5, 9.

Hunter, P., Wallace, D., 2001: Lumpy skin disease in southern Africa: a review of the disease and aspects of control. J S Afr Vet Assoc 72:68–71

Jarvis, A., Reuter, H.I., Nelson, A., Guevara, E., 2008. Hole-filled SRTM for the globe version 4. Available from CGIAR-CSI SRTM 90 m database. https://doi.org/10.1055/s-0029-1222523

Hopp, M.J., Foley, J.A., 2001. Global-scale relationships between climate and the dengue fever vector, AEDES AEGYPTI. Clim. Change. https://doi.org/10.1023/A:1010717502442

Kahana-Sutin, E., Klement, E., Lensky, I., Gottlieb, Y., 2017. High relative abundance of the stable fly Stomoxys calcitrans is associated with lumpy skin disease outbreaks in Israeli dairy farms. Med. Vet. Entomol. https://doi.org/10.1111/mve.12217

Klausner, Z., Fattal, E., Klement, E., 2017. Using Synoptic Systems’ Typical Wind Trajectories for the Analysis of Potential Atmospheric Long-Distance Dispersal of Lumpy Skin Disease Virus. Transbound. Emerg. Dis. https://doi.org/10.1111/tbed.12378

Knorr-Held, L., Besag, J., 1998. Modelling risk from a disease in time and space. Stat. Med. https://doi.org/10.1002/(SICI)1097-0258(19980930)17:18<2045::AID-SIM943>3.0.CO;2-P

Lawson, A. B. 2013. Bayesian Disease Mapping: Hierarchical Modeling in Spatial Epidemiology. New York, CRC Press

Lawson, A.B., 2018. Bayesian latent modeling of spatio-temporal variation in small-area health data. Wiley Interdiscip. Rev. Comput. Stat. https://doi.org/10.1002/wics.1441

Lesaffre, E., Lawson, A.B., 2012. Bayesian Biostatistics, Bayesian Biostatistics. https://doi.org/10.1002/9781119942412

Zehang, L., Yuan, H., Godwin, J., Martin, D. M., Wakefield, J., Clark, S. J., with support from the United Nations Inter-agency Group for Child Mortality Estimation and its technical advisory group. 2019. Changes in the spatial distribution of the under-five mortality rate: Small-area analysis of 122 DHS surveys in 262 subregions of 35 countries in Africa. PLoS One. 14(1): e0210645. https://doi.org/10.1371/journal.pone.0210645

Lubinga, J.C., Tuppurainen, E.S.M., Mahlare, R., Coetzer, J.A.W., Stoltsz, W.H., Venter, E.H., 2015. Evidence of transstadial and mechanical transmission of lumpy skin disease virus by Amblyomma hebraeum ticks. Transbound. Emerg. Dis. https://doi.org/10.1111/tbed.12102

Mercier, A., Arsevska, E., Bournez, L., Bronner, A., Calavas, D., Cauchard, J., Falala, S., Caufour, P., Tisseuil, C., Lefrançois, T., Lancelot, R., 2018. Spread rate of lumpy skin disease in the Balkans, 2015–2016. Transbound. Emerg. Dis. https://doi.org/10.1111/tbed.12624

Merow, C., Smith, M.J., Silander, J.A., 2013. A practical guide to MaxEnt for modeling species’ distributions: What it does, and why inputs and settings matter. Ecography (Cop.). https://doi.org/10.1111/j.1600-0587.2013.07872.x

Muscarella, R., Galante, P.J., Soley-Guardia, M., Boria, R.A., Kass, J.M., Uriarte, M., Anderson, R.P., 2014. ENMeval: An R package for conducting spatially independent evaluations and estimating optimal model complexity for Maxent ecological niche models. Methods Ecol. Evol. https://doi.org/10.1111/2041-210X.12261

Ochwo, S., VanderWaal, K., Munsey, A., Ndekezi, C., Mwebe, R., Okurut, A.R.A., Nantima, N., Mwiine, F.N. 2018. Spatial and temporal distribution of lumpy skin disease outbreaks in Uganda (2002-2016). BMC Vet. Res. https://doi.org/10.1186/s12917-018-1503-3

OIE (2016) Lumpy skin disease. In: OIE Terrestrial Animal Health Code. http://web.oie.int/eng/normes/mcode/en_chapitre_1.11.12.htm#rubrique_dermatose_nodulaire_contagieuse. Accessed 30 Nov 2018.

OIE. 2018a. Lumpy Skin Disease in: Technical disease cards. URL: http://www.oie.int/fileadmin/Home/eng/Animal_Health_in_the_World/docs/pdf/Disease_cards/LUMPY_SKIN_DISEASE_FINAL.pdf (accessed 17 December 2018)

OIE. 2018b. OIE-Listed diseases, infections and infestations in force in 2018. URL: http://www.oie.int/animal-health-in-the-world/oie-listed-diseases-2018/ (accessed 17 December 2018)

OIE, WAHID. 2018. World Animal Health Information Database. URL: http://www.oie.int/wahis_2/public/wahid.php/Wahidhome/Home. accessed 17 December 2018

Patz, J.A., Campbell-Lendrum, D., Holloway, T., Foley, J.A., 2005. Impact of regional climate change on human health. Nature. https://doi.org/10.1038/nature04188

Peterson, A.T., Papeş, M., Soberón, J., 2008. Rethinking receiver operating characteristic analysis applications in ecological niche modeling. Ecol. Modell. https://doi.org/10.1016/j.ecolmodel.2007.11.008

Phillips, S.J., Anderson, R.P., Schapire, R.E., 2006. Maximum entropy modeling of species geographic distributions. Ecol. Modell. https://doi.org/10.1016/j.ecolmodel.2005.03.026

Qiao, H., Soberón, J., Peterson, A.T., 2015. No silver bullets in correlative ecological niche modelling: Insights from testing among many potential algorithms for niche estimation. Methods Ecol. Evol. https://doi.org/10.1111/2041-210X.12397

Radosavljevic, A., Anderson, R.P., 2014. Making better Maxent models of species distributions: Complexity, overfitting and evaluation. J. Biogeogr. https://doi.org/10.1111/jbi.12227

Richardson, S., Thomson, A., Best, N., Elliott, P., 2004. Interpreting posterior relative risk estimates in disease-mapping studies. Environ. Health Perspect. https://doi.org/10.1289/ehp.6740

Riebler, A., Sørbye, S.H., Simpson, D., Rue, H., Lawson, A.B., Lee, D., MacNab, Y., 2016. An intuitive Bayesian spatial model for disease mapping that accounts for scaling, in: Statistical Methods in Medical Research. https://doi.org/10.1177/0962280216660421

Robinson, T.P., William Wint, G.R., Conchedda, G., Van Boeckel, T.P., Ercoli, V., Palamara, E., Cinardi, G., D’Aietti, L., Hay, S.I., Gilbert, M., 2014. Mapping the global distribution of livestock. PLoS One. https://doi.org/10.1371/journal.pone.0096084

Romero-Alvarez, D., Escobar, L.E., Varela, S., Larkin, D.J., Phelps, N.B.D., 2017. Forecasting distributions of an aquatic invasive species (Nitellopsis obtusa) under future climate scenarios. PLoS One. https://doi.org/10.1371/journal.pone.0180930

Rue, H., Martino, S., Chopin, N., 2009. Approximate Bayesian inference for latent Gaussian models by using integrated nested Laplace approximations. J. R. Stat. Soc. Ser. B Stat. Methodol. https://doi.org/10.1111/j.1467-9868.2008.00700.x

Saegerman, C., Bertagnoli, S., Meyer, G., Ganière, J.P., Caufour, P., De Clercq, K., Jacquiet, P., Fournie, G., Hautefeuille, C., Etore, F., Casal, J., 2018. Risk of introduction of lumpy skin disease in France by the import of vectors in animal trucks. PLoS One. https://doi.org/10.1371/journal.pone.0198506

Simpson, D., Rue, H., Riebler, A., Martins, T. G. & Sorbye, S. H. 2017. Penalising Model Component Complexity: A Principled, Practical Approach to Constructing Priors. Statistical Science 32, 1–28

Soberón, J., Peterson, A.T., 2005. Interpretation of Models of Fundamental Ecological Niches and Species’ Distributional Areas. Biodivers. Informatics. https://doi.org/10.17161/bi.v2i0.4

Spiegelhalter, D.J., Best, N.G., Carlin, B.P., Van Der Linde, A., 2002. Bayesian measures of model complexity and fit. J. R. Stat. Soc. Ser. B Stat. Methodol. https://doi.org/10.1111/1467-9868.00353

Tageldin, M.H., Wallace, D.B., Gerdes, G.H., Putterill, J.F., Greyling, R.R., Phosiwa, M.N., Al Busaidy, R.M., Al Ismaaily, S.I., 2014. Lumpy skin disease of cattle: An emerging problem in the Sultanate of Oman. Trop. Anim. Health Prod. https://doi.org/10.1007/s11250-013-0483-3

Tasioudi, K.E., Antoniou, S.E, Iliadou, P., Sachpatzidis, A., Plevraki, E., Agianniotaki, E.I., Fouki, C., Mangana-Vougiouka, O., Chondrokouki, E., Dile, C., 2016. Emergence of lumpy skin disease in Greece, 2015. Transbound Emerg Dis 63:260–265

Terrestrial Animal Health Code. 2018. OIE. Chapter 11.9. Infection with Lumpy Skin Disease Virus. Available online at http://www.oie.int/index.php?id=169&L=0&htmfile=chapitre_lsd.htm (accessed 17 December 2018)

Tuppurainen, E.S.M., Oura, C.A.L., 2012. Review: Lumpy Skin Disease: An Emerging Threat to Europe, the Middle East and Asia. Transbound. Emerg. Dis. https://doi.org/10.1111/j.1865-1682.2011.01242.x

Tuppurainen, E.S.M., Stoltsz, W.H., Troskie, M., Wallace, D.B., Oura, C.A.L., Mellor, P.S., Coetzer, J.A.W., Venter, E.H., 2011. A Potential Role for Ixodid (Hard) Tick Vectors in the Transmission of Lumpy Skin Disease Virus in Cattle. Transbound. Emerg. Dis. https://doi.org/10.1111/j.1865-1682.2010.01184.x

Tuppurainen, E.S.M., Lubinga, J.C., Stoltsz, W.H., Troskie, M., Carpenter, S.T., Coetzer, J.A.W., Venter, E.H., Oura, C.A.L., 2013. Evidence of vertical transmission of lumpy skin disease virus in Rhipicephalus decoloratus ticks. Ticks Tick. Borne. Dis. https://doi.org/10.1016/j.ttbdis.2013.01.006

Tuppurainen, E.S.M., Babiuk, S., Klement, E. 2018. Lumpy Skin Disease. VI, 109. Springer Int Publishing. DOI: 10.1007/978-3-319-92411-3

Tuppurainen, E. S. M., Antoniou, S. E., Tsiamadis, E., Topkaridou, M., Labus, T., Debeljak, Z6., Plavšić, B., Miteva, A., Alexandrov, T., Pite, L., Boci, J., Marojevic, D., Kondratenko, V1., Atanasov, Z., Murati, B., Acinger-Rogic, Z., Kohnle, L., Calistri, P., Broglia, A. 2018. Field observations and experiences gained from the implementation of control measures against lumpy skin disease in South-East Europe between 2015 and 2017. In press, https://doi.org/10.1016/j.prevetmed.2018.12.006.

Warren, D.L., Seifert, S.N., 2011. Ecological niche modeling in Maxent: The importance of model complexity and the performance of model selection criteria. Ecol. Appl. https://doi.org/10.1890/10-1171.1

Wickham, H., 2016. tidyverse: Easily Install and Load “Tidyverse” Packages., R package version 1.0.0. https://doi.org/10.1016/j.smallrumres.2011.03.043

Woods, J.A., 1988. Lumpy skin disease-A review. Trop. Anim. Health Prod. https://doi.org/10.1007/BF02239636

