## Supplementary figures and images for "Mapping changes in the spatiotemporal distribution of lumpy skin disease virus"

### Fitted RR spacextime

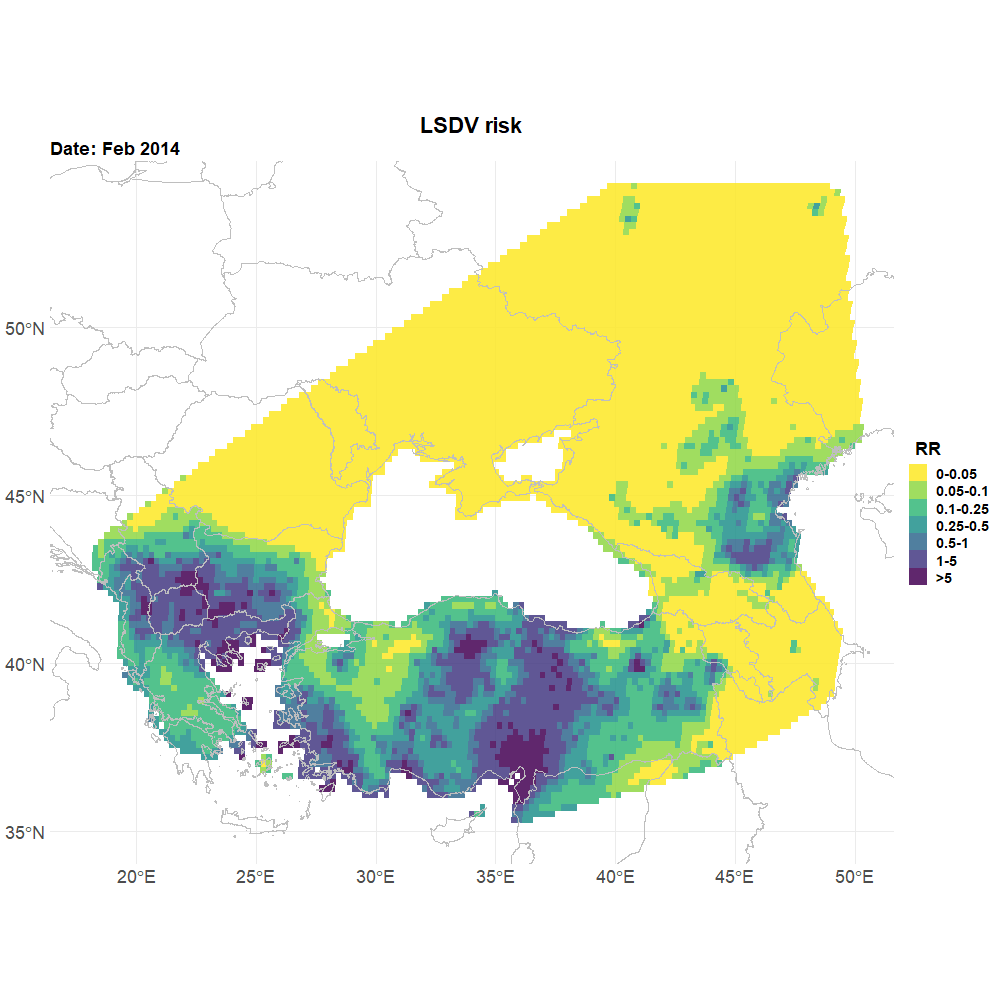
